# Reciprocal repression between Isl1 and Nkx2-5 defines the zebrafish embryonic cardiac pacemaker domain

**DOI:** 10.64898/2026.10.16.759628

**Authors:** F. Tessadori, L. Garric, L. Blankenborg, A. Kisoensingh, R.R. Padala, T.P. de Boer, J. Bakkers

## Abstract

The cardiac pacemaker initiates each heartbeat and is essential for normal cardiac function. Although the gene regulatory network controlling pacemaker development has been extensively studied in mammals, it remains unclear whether the same regulatory hierarchy governs pacemaker development in other vertebrates. Here, we combined high-speed optical mapping with genetic analysis to define the gene regulatory network that establishes and maintains pacemaker identity in the zebrafish embryo. We show that the LIM-homeodomain transcription factor Isl1 is essential for maintaining pacemaker identity of venous pole cardiomyocytes. Loss of Isl1 causes pacemaker cells to acquire chamber-like characteristics through ectopic *nkx2-5* expression, resulting in loss of a dominant pacemaker site and severe bradyarrhythmia. Reducing Nkx2-5 activity largely restores regular rhythm despite persistent loss of other Isl1-dependent markers, indicating that inappropriate Nkx2-5 activity contributes substantially to the electrophysiological defects in *isl1* mutants. In contrast, genetic disruption of *shox2* does not impair embryonic pacemaker development or alter *isl1* expression, suggesting that Shox2 is not an essential component of the zebrafish pacemaker gene regulatory network. Together, our findings establish a zebrafish-specific transcriptional hierarchy in which Isl1 safeguards pacemaker identity by restricting Nkx2-5 independently of Shox2. These results show that a conserved boundary between pacemaker and chamber myocardium can be maintained by distinct upstream factors during vertebrate heart development.

## Introduction

The rhythmic contraction of the vertebrate heart relies on a specialized population of pacemaker cells located in the sinoatrial node (SAN) in mammals and at the venous pole of the zebrafish heart (Burkhard et al., 2017; Martin and Waxman, 2021; van der Maarel and Christoffels, 2024; van Weerd and Christoffels, 2016). Unlike working atrial and ventricle cardiomyocytes, pacemaker cells generate spontaneous action potentials that initiate each heartbeat. The establishment of a functional pacemaker therefore requires not only the specification of pacemaker cells during embryogenesis, but also the maintenance of a distinct pacemaker identity while neighboring cardiomyocytes differentiate into chamber myocardium. Although the molecular mechanisms underlying pacemaker specification have been studied extensively, (reviewed in Park and Fishman, 2026; van Eif et al., 2018) considerably less is known about how pacemaker identity is maintained once these cells have differentiated.

Cardiac pacemaker cells originate from the second heart field progenitors and are molecularly distinct from those giving rise to chamber cardiomyocytes. Recent work demonstrated that antagonistic BMP and Hedgehog signaling establish the inflow tract progenitor field early in zebrafish development, thereby defining the future pacemaker domain (Robertson et al., 2026). As these progenitors differentiate, pacemaker and chamber cardiomyocytes acquire distinct transcriptional identities. In zebrafish, pacemaker cells co-express *isl1* with *hcn4*, *bmp4* and *tbx2b*, while the adjacent chamber cardiomyocytes express *nkx2-5* and activate the chamber myocardial gene program (Colombo et al., 2018; Targoff et al., 2013; Tessadori et al., 2012). Studies in mouse have identified *Isl1, Shox2, Tbx3, Tbx18* and *Nkx2-5* as central components of the pacemaker gene regulatory network. *Tbx3, Shox2* and *Isl1* drive the pacemaker gene program, while *Nkx2-5* specifies and maintains chamber myocardial identity, suppressing pacemaker genes in the differentiating chamber myocardium (Blaschke et al., 2007; Cai et al., 2003; Christoffels et al., 2006; de Pater et al., 2009; Liang et al., 2015; Mommersteeg et al., 2007). Together this indicates a conserved process in which early signalling pathways specify the pacemaker progenitor population, after which transcriptional regulators maintain the boundary between pacemaker and chamber myocardium.

As pacemaker differentiation proceeds, the mammalian SAN remains NKX2-5-negative due to the repressive activity of homeodomain transcription factor SHOX2 (Espinoza-Lewis et al., 2009), preventing their differentiation into chamber myocardium (Blaschke et al., 2007; Espinoza-Lewis et al., 2009). Isl1 has a distinct role within this network. In addition to its early function in second heart field development, Isl1 remains expressed in differentiated pacemaker cells throughout life. Conventional *Isl1*-null mouse embryos fail to form multiple second heart field-derived structures, including the SAN itself, precluding analysis of later pacemaker functions (Cai et al., 2003). Conditional deletion of *Isl1* in *Hcn4*+ CMs results in reduced heart rate (Liang et al., 2015) and is accompanied by reduction of expression of pacemaker genes *Hcn4, Shox2* and *Tbx3*. However, these mice remain viable and, unlike *Shox2* or *Tbx3* deficiency, *Isl1* deletion does not lead to ectopic *Nkx2-5* expression within the SAN (Liang et al., 2015). These findings indicate that ISL1 is required for the pacemaker gene program, but it remains unclear whether it actively maintains pacemaker identity or acts indirectly by preventing repression of pacemaker genes.

The zebrafish model offers a unique opportunity to dissect these developmental functions. In contrast to mice, loss of *isl1* function does not prevent formation of second heart field derivatives or incorporation of venous pole cardiomyocytes in the heart (Burkhard and Bakkers, 2018). Instead, *isl1* mutants develop a characteristic and lethal bradyarrhythmia phenotype (de Pater et al., 2009; Tessadori et al., 2012), allowing the function of Isl1 to be investigated after pacemaker cells have formed, providing a model to investigate how pacemaker identity is maintained. Here, we combine high-speed optical voltage imaging with genetic perturbation and molecular analysis to define the gene regulatory network that maintains pacemaker identity in the embryonic zebrafish heart. We show that loss of Isl1 disrupts stable action potential initiation at the venous pole, resulting in erratic pacemaker activity and impaired cardiac rhythm. Unexpectedly, *shox2*/Shox2 is dispensable for pacemaker function in zebrafish. Instead, Isl1 maintains pacemaker identity by preventing *nkx2-5* expression in the pacemaker domain, and partial loss of Nkx2-5 restores regular rhythm in *isl1* mutants. Together, our findings identify reciprocal repression between Isl1 and Nkx2-5 as a transcriptional circuit that stabilizes pacemaker identity after its initial specification and preserves the functional boundary between pacemaker and chamber myocardium; and indicate an evolutionary divergence in the network with regard to the requirement of Isl1 (zebrafish) and Shox2 (mouse) in the suppression of the conserved chamber boundary function of Nkx2-5.

## Results

### Isl1 establishes a dominant site of action potential initiation at the venous pole

To determine how Isl1 regulates embryonic pacemaker function, we analyzed electrical activity using the myocardial-specific transgenic FRET voltage reporter *tg(myl7:chimeric VSFP-butterfly CY)* (van Opbergen et al., 2018), referred to as VSFP-butterfly fish in the rest of this study. To eliminate motion artefacts, cardiac contraction was stopped by injecting a *tnnt2a* morpholino at the 1-cell stage (Sehnert et al., 2002), allowing high-speed optical recordings of action potentials at 3 days post fertilization (dpf) (Figure 1A, left panel). Ratiometric voltage sensor signals were depicted in spatiotemporal maps from which the site of the action potential (APs) initiation, propagation velocity and repolarization kinetics could be quantified. To distinguish pacemaker-specific defects from more general abnormalities in cardiac electrophysiology, we compared Isl1-deficient embryos to Tbx5a-deficient embryos Figure 1B). F0 CRISPR/Cas9 Crispants faithfully reproduced the corresponding deficiency phenotypes (de Pater et al., 2009; Garrity et al., 2002; Tessadori et al., 2021; Tessadori et al., 2012) and were therefore used throughout the functional analyses (Supplementary Figure S1). We first examined repolarization at 3 dpf by measuring APD_50_. Loss of Tbx5a prolongated APD_50_ in both atrium and ventricle, whereas Isl1 deficiency affected ventricular repolarization only (Figure 1C, D) suggesting the absence of Isl1 does not affect the repolarization capacity of the venous pole CMs. Next, we quantified AP propagation from the atrium to the ventricle by measuring the time (dt) between arrival of depolarization peaks at both sites (Figure 1E). Although AP conduction was significantly slower in *tbx5a* crispants the average conduction time was unchanged in *isl1* crispants (Figure 1F). Instead, *isl1* crispants displayed marked beat-to-beat variability in conduction intervals (Figure 1G), suggesting that AP initiation rather than propagation was affected. To directly test this possibility, we mapped the site where the first AP of each beat was initiated along the cardiac posterior-anterior axis (Figure 1H). In wild-type and *tbx5a* crispants, a majority of the APs originated from the venous pole. In contrast, AP initiation in *isl1* crispants displayed a more spread distribution, indicating that AP initiation events occurred at multiple locations throughout the atrium, demonstrating loss of a dominant pacemaker site, and consequently unmasking residual automaticity of zebrafish CMs at 3 dpf. Such ectopic initiation is consistent with the intrinsic automaticity of early heart tube cardiomyocytes, which is detectable when the fastest pacemaker (ie the Isl1-positive domain at the venous pole here) is lost or suppressed (Van Mierop, 1967). Together, these findings demonstrate that while Tbx5a is required for fast conduction of the AP through the atrium and ventricle, Isl1 is dispensable for propagation of the AP through the atrium, but is essential for establishing the dominance of the venous pole as the site of pacemaker activity during zebrafish heart development.

**Figure 1:**
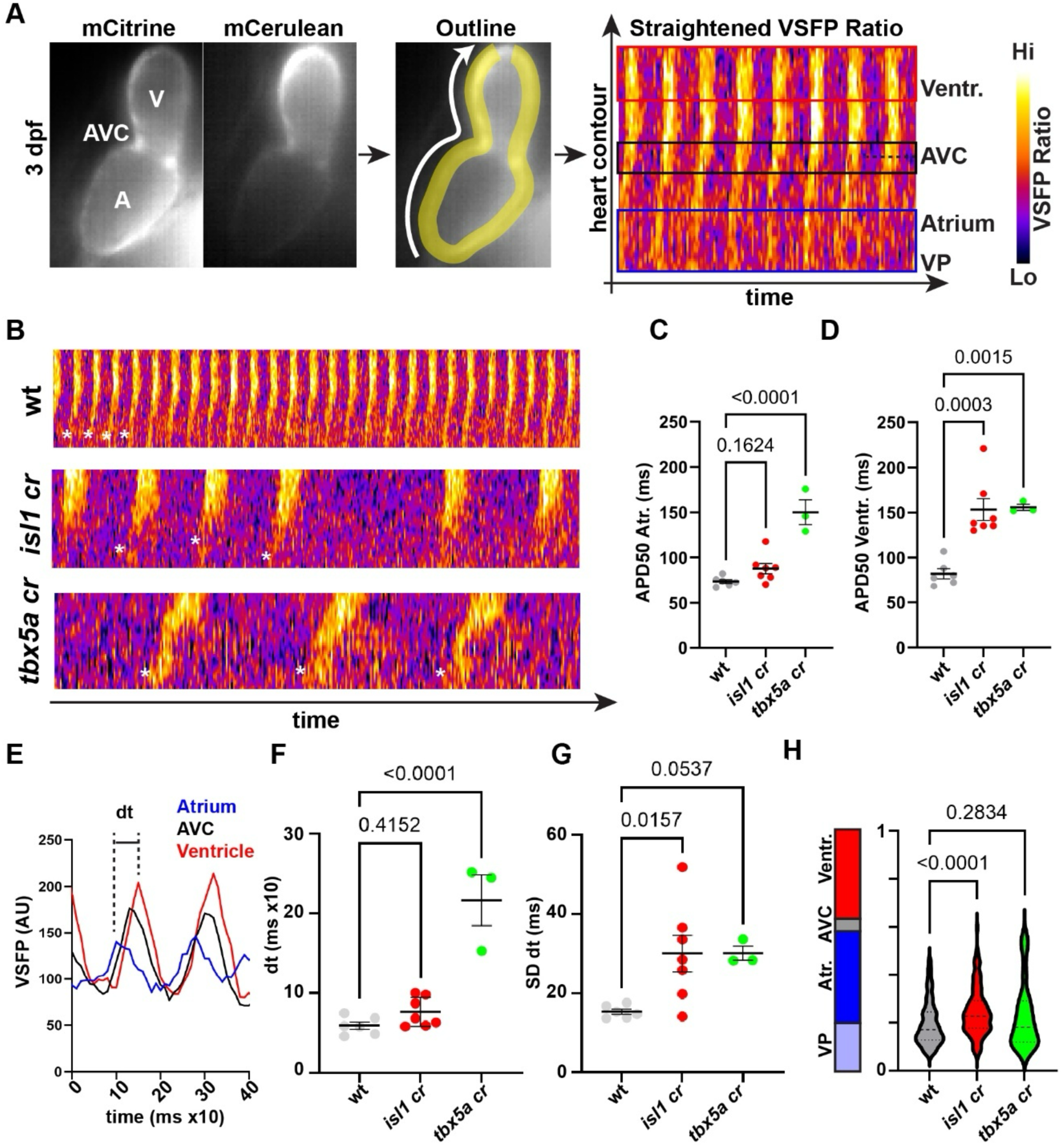
In vivo recordings of APs reveal loss of dominance of the venous pole in Isl1-defective embryos. **(A)** From left to right: illustrative stills of the mCitrine and mCerulean recordings carried out on 3 dpf *tg(myl7:VSFP-CY-Butterfly)* larvae. After image processing (see Materials and Methods section and van Opbergen et al., 2018), ratiometric heatmaps of fluorescent intensities are extracted that reveal AP propagation features in the different segments of the heart. Boxes are illustrative for the profiles presented in panel (E). **(B)** Comparison of AP profiles in wt, Isl1-deficient and Tbx5a deficient backgrounds. The difference in overall profile shape denotes between the panels illustrates difference in AP propagation speed, repolarization and conduction properties of the heart segments. White asterisks illustrate the detected start of the AP for the corresponding profile. **(C)** Atrial APD50 in wt, *isl1* and *tbx5a* crispants. **(D)** Ventricular APD50 calculated in the same recordings as presented in (C) **(E)** Illustrative profiles extracted from AP heatmaps presented in panel (A) for different segments of the heart. Calculation of average dt and SD dt as presented in panels (F) and (G) is based on peaks obtained on the Atrium and Ventricle profiles. **(F)** Average dt in wt, *isl1* crispant *tbx5a* crispant larvae. **(G)** SD dt for the same dataset as dt reported in (F). **(H)** Mapping of AP points of origin (see asterisks in panel (B)). For all panels, one-way ANOVA with multiple comparisons, wt values taken as reference. Bars are at mean +/-SEM.

### Loss of Isl1 promotes chamber myocardial identity in venous pole cardiomyocytes

Having established that Isl1 is required for stable pacemaker activity, we next asked whether Isl1 also maintains the molecular identity of pacemaker cells. At 52 hpf, when the arrhythmic phenotype first becomes apparent in *isl1* mutants, the venous pole is characterized by expression of pacemaker-associated genes including *bmp4, shox2, hcn4* and *tbx2b*, whereas the adjacent atrial myocardium expresses *nppa* (Burkhard and Bakkers, 2018; Colombo et al., 2018; de Pater et al., 2009). Whole mount in situ hybridization (ISH; Figure 2) revealed a selective loss of pacemaker gene expression at the venous pole in *isl1^-/-^* hearts. Expression of *tbx2b* and *bmp4* was abolished specifically at the venous pole, while remaining unchanged at the atrioventricular canal, demonstrating that Isl1 is specifically required to maintain the pacemaker domain rather than non-working myocardium generally. Likewise, expression of *hcn4,* which encodes the hyperpolarization-activated potassium channel underlying the spontaneous pacemaker depolarization (Baruscotti et al., 2011; Colombo et al., 2018; Shi et al., 1999; Tessadori et al., 2012; Yamamoto et al., 2006), was lost from the venous pole but remained detectable in the AVC and inner curvature of the ventricle. In contrast, the chamber myocardium marker *nppa* expanded towards the inflow tract suggesting that venous pole cardiomyocytes acquire a chamber-like identity in the absence of Isl1. We also observed a complete loss of *shox2* from the venous pole in *isl1^-/-^* hearts. In mammals, Shox2 controls Isl1 expression to maintain sinoatrial node identity (Hoffmann et al., 2013). The dependence of *shox*2 expression on Isl1 therefore suggests a distinct regulatory relationship in zebrafish and raises the possibility that *isl1* acts upstream of *shox2.* Together, these findings demonstrate that Isl1 is required to maintain the molecular identity of venous pole pacemaker cells. Loss of Isl1 results in coordinated downregulation of the pacemaker transcriptional program accompanied by expansion of chamber myocardial identity, consistent with the functional defects described above.

**Figure 2:**
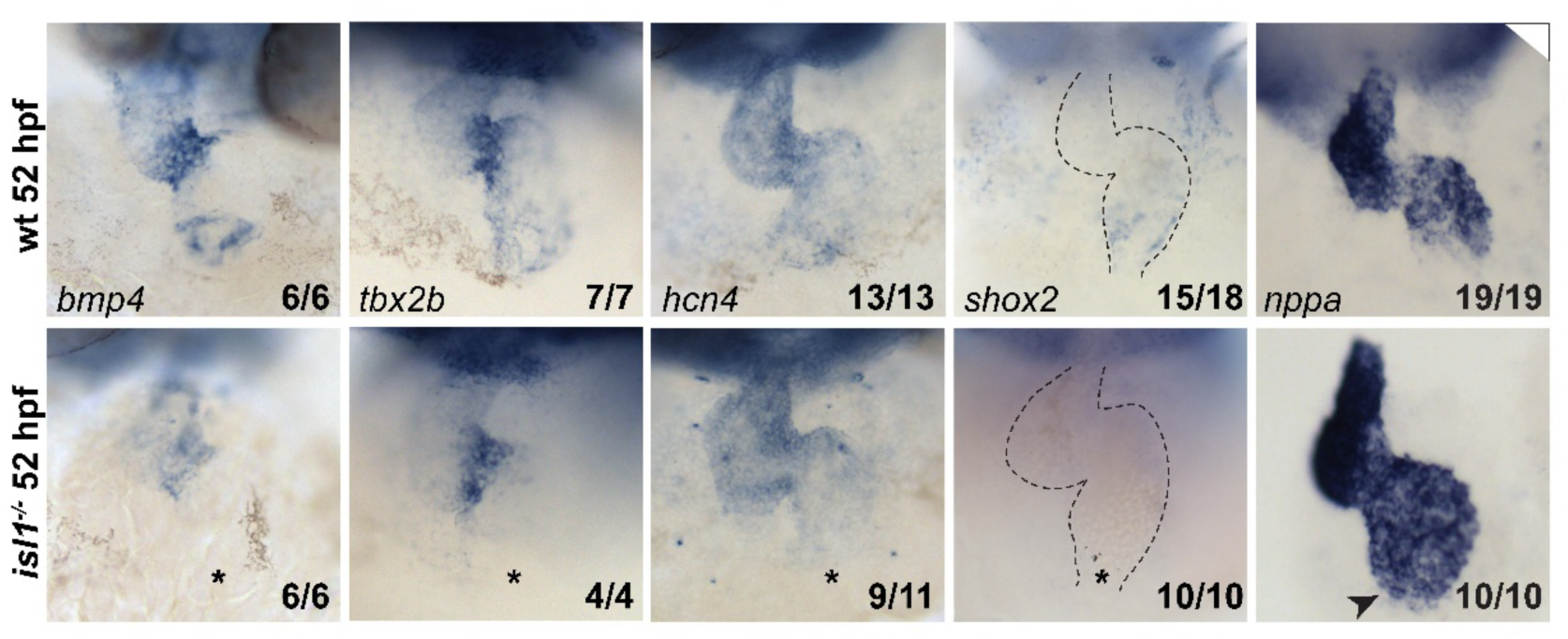
Gene patterning is affected at the venous pole of *isl1^-/-^* embryos. Whole mount RNA is situ hybridization on 52 hpf embryos. Expression of *bmp4, tbx2b*, *hcn4* and *shox2* is lost at the VP of *isl1^-/-^* embryos (asterisk), but remains unaffected in other expressing regions of the heart (AVC and/or inner curvature of the ventricle for *bmp4*, *tbx2b* and *hcn4*). Expression of *shox2*, normally present only at the VP in the wt, is lost in *isl1^-/-^* mutants. The domain of expression of *nppa* in the atrium appears expanded towards the VP (arrowhead) in *isl1^-/-^* embryos. Number of embryos displaying the presented phenotype / total number of analyzed embryos is presented in the bottom-right corner of each panel.

### Independent *shox2* mutant alleles demonstrate Shox2 is dispensable for embryonic pacemaker development

Because *shox2* expression is lost in *isl1* mutants, we next asked whether Shox2 is required for embryonic pacemaker development in zebrafish. Previous morpholino-mediated knockdown studies in zebrafish suggested that *shox*2 is required for normal pacemaker function by regulating *isl1* expression (Hoffmann et al., 2013), but the genetic requirement for Shox2 has remained unresolved. We therefore generated 3 different mutant alleles of *shox2* predicted to disrupt Shox2 function through distinct mechanisms: an RNA-less allele (*shox2_RL),* a truncation of the protein upstream of the homeodomain (*shox2_HD*) and a C-terminal truncation (*shox2_TR*) (Figure 3A). In *shox2_RL*^-/-^ embryos we observed marked reduction of *shox2* expression at 26 hpf and 52 hpf in cephalic structures where it is normally strongly present and required for neuronal development, indicating efficient disruption of gene expression (Figure 3B). Both homozygous *shox2_TR* and *shox2_RL* larvae showed reduced survivability beyond the larval stage (approximately 9% and 2% at 10 weeks, respectively; data not shown), likely due to failure of the larvae to inflate the swim bladder, which is consistent with the phenotype reported in a previously published *shox2* null allele (Laureano et al., 2022), suggesting that our mutant alleles suppress Shox2 function.

**Figure 3:**
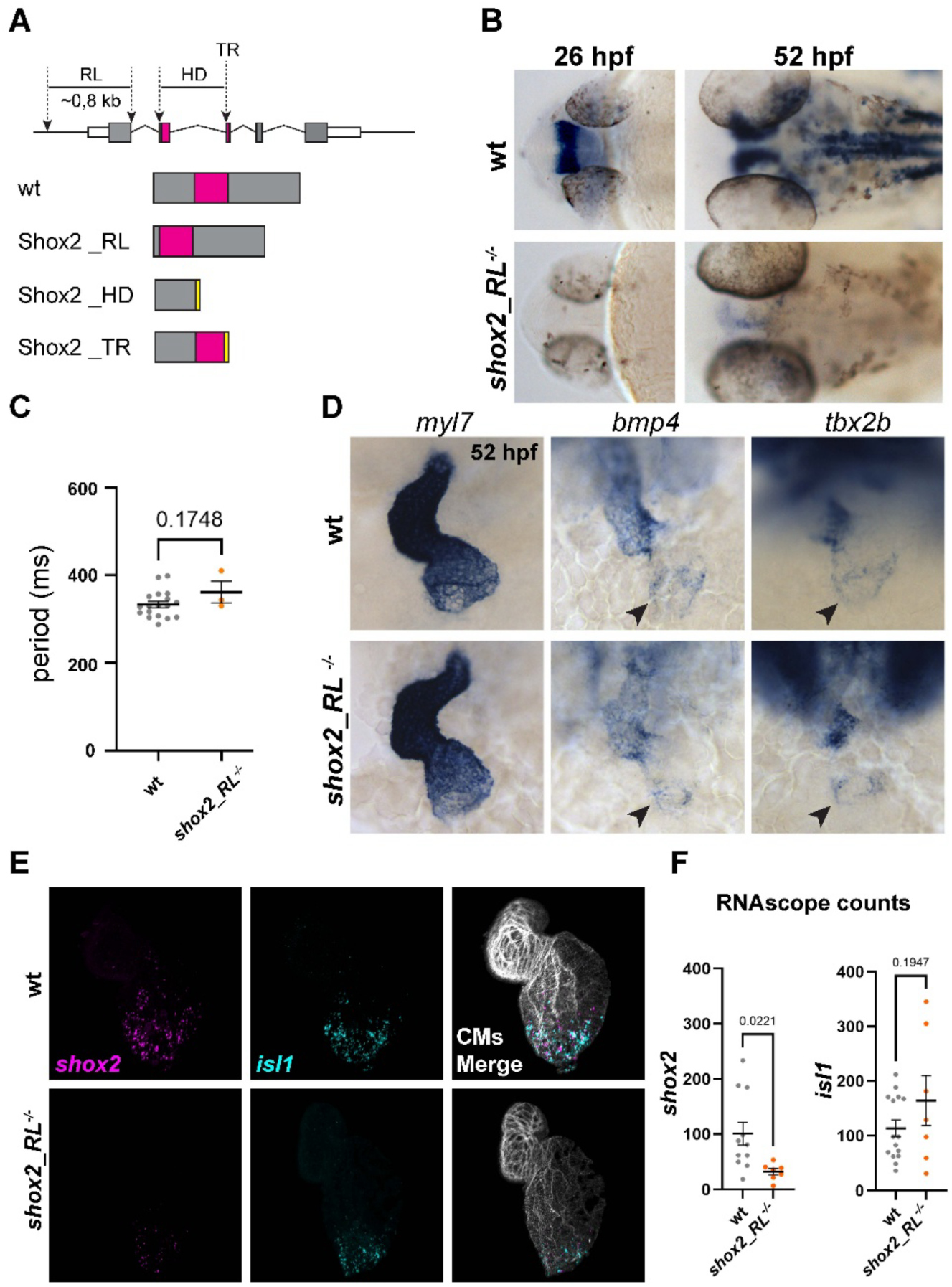
Mutations in *shox2* do not affect *isl1* expression and pacemaker function in zebrafish embryos. **(A)** The *shox2* locus genomic deletions (dashed arrows) and predicted translation products for the shox2 alleles presented in this study. RL: RNAless, TR: truncation, HD: homeodomain. All data in the additional panels of the figure are obtained on the Shox2_RL line **(B)** Whole mount RNA ISH at 26 hpf and 52 hpf displays virtually full loss of *shox2* expression in the *shox2_RL^-/-^* embryos. **(C)** Heartbeat period of *shox2_RL^-/-^* larvae and corresponding wt embryos at 5 dpf. **(D)** *Shox2_RL^-/-^*embryos show apparent normal cardiac morphogenesis (*myl7*) and normal expression pattern of AVC and VP markers *bmp4* and *tbx2b*. Arrowheads indicate position of the VP. **(E)** Double RNAscope on *shox2* (magenta) and *isl1* (cyan) at 52 hpf reveals sustained expression of *isl1* at the cardiac VP in a Shox2-defective genetic background. **(F)** Quantification of RNAscope counts as detected in (E). Two-tailed, non-paired Student’s t-test; p value is reported in the figure panel. Bars are at mean +/-SEM.

We next examined whether disruption of Shox2 affects *isl1* expression. Unexpectedly *isl1* expression remained unchanged in *shox2_RL^-/^*^-^ mutant hearts (Figure 3E, F), indicating that reduced *shox2* expression does not obviously affect maintenance of *isl1*.

Furthermore, homozygous mutants for all three alleles exhibited heart rates indistinguishable from wild-type siblings a 5 dpf (Figure 3C and Supplementary Figure S3A). Likewise, cardiac morphology appeared normal, as assessed by ISH for the myocardial marker *myl7*, and the venous pole markers *bmp4* and *tbx2b* retained their normal expression domains, indicating that specification of the pacemaker region was not detectably affected (Figure 3D). One possible explanation for the absence of a cardiac phenotype is genetic compensation (El-Brolosy et al., 2019) by the closely related gene *shox* (Kenyon et al., 2011; Rao et al., 1997). However, overall *shox* expression levels remained unaffected in any of our *shox2* homozygous alleles and, specifically, absent from the VP (Supplementary Figure S3B, C).

Overall, these analyses failed to reveal detectable defects in embryonic pacemaker specification, molecular patterning or cardiac rhythm in any of the three independently generated *shox2* mutant alleles. These findings contrast with previous morpholino-based studies and indicate that severe genetic disruption of *shox2* is not sufficient to impair embryonic pacemaker development or maintenance of *isl1* expression.

### Isl1 maintains pacemaker identity by excluding Nkx2-5 from the venous pole

Our analysis of *shox2* mutant embryos did not reveal detectable defects in pacemaker development, suggesting that Isl1 maintains pacemaker identity through a Shox2-independent mechanism. In mammals, repression of Nkx2-5 is essential for sinoatrial node specification (Espinoza-Lewis et al., 2009). We therefore asked whether Isl1 restricts the Nkx2-5 expression domain during zebrafish heart development.

At the cardiac disc stage (22 somites), Nkx2-5 and *isl1* occupy complementary expression domains (Figure 4A; Ren et al., 2019), with Nkx2-5 expression in the inner diameter of the cardiac disc and *isl1* expression at the periphery of the disc. This spatial segregation is maintained during cardiac jogging, when *isl1* becomes restricted to a ring of cells at the venous pole, immediately adjacent to the Nkx2-5-positive atrium (Figure 4B). By 52 hpf, the pacemaker domain remains devoid of detectable Nkx2-5 expression (Figure 4C).

**Figure 4:**
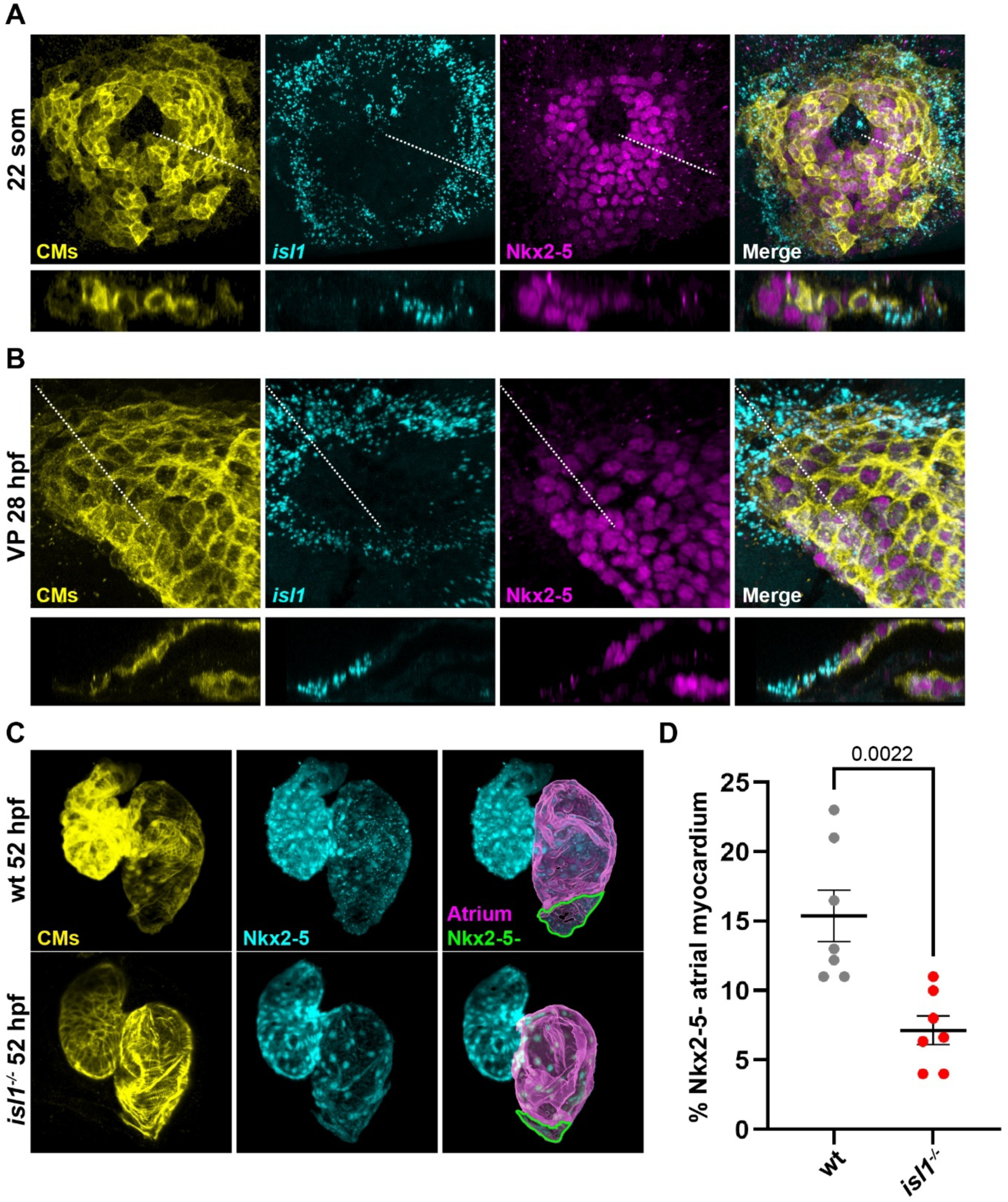
Loss of Isl1 shifts the boundary of the Nkx2-5 expression domain in the embryonic zebrafish heart. **(A)** Combination of RNAscope (*isl1;* cyan) and fluorescence immunolabeling (CMs, yellow; Nkx2-5, magenta) shows that at 22 somites the domains of expression of *isl1* and Nkx2-5 are mutually exclusive in the cardiac disc. The lower panels are optical sections of the corresponding higher panels at the dashed line. **(B)** At the end of cardiac jogging the *isl1* expression domain forms a ring at the venous pole of the heart tube. Nkx2-5 expression is present in the atrium but does not overlap with *isl1*+ cells. The lower panels are optical sections of the corresponding higher panels at the dashed line. **(C)** Fluorescent immunolabeling of CMs (yellow) and Nkx2-5 (cyan) in wt and *isl1^-/-^* embryos at 52 hpf. Atria are overlayed in magenta and the Nkx2-5-VP is outlined in green. **(D)** Quantification of the Nkx2-5-atrial myocardium obtained on confocal imaging as presented in panel **(C)**. Two-tailed, non-paired Student’s t-test; p value is reported in the figure panel. Bars are at mean +/- SEM.

To determine whether Isl1 establishes this boundary, we examined Nkx2-5 expression in *isl1* mutant embryos. In wild-type hearts, the Nkx2-5-negative venous pole comprises approximately 15% of the atrial myocardium (Figure 4C, D). In contrast, this domain was reduced by more than half in *isl1* mutants owing to expansion of Nkx2-5 expression towards the venous pole. Thus, although venous pole cardiomyocytes are specified and incorporated into the heart in the absence of Isl1, they fail to maintain a molecular pacemaker identity and instead acquire features of chamber myocardium. To determine whether this regulation depends on Shox2, we quantified the Nkx2-5-negative pacemaker domain in *shox2_RL^-/-^* embryos. In contrast to *isl1* mutants, *shox2* mutants exhibited a normal-sized Nkx2-5-negative VP (Supplementary Figure S4A, B), indicating that restriction of Nkx2-5 to the atrium does not depend on detectable *shox2* expression.

Collectively, these findings indicate that Isl1 is required to restrict Nkx2-5 expression at the venous pole, while disruption of Shox2 did not measurably alter the Nkx2-5 expression domain.

### Reducing Nkx2-5 activity restores regular pacing in isl1-deficient embryos

Our data suggested that the pacemaker defects observed in *isl1* mutants result from ectopic expansion of Nkx2-5 into the venous pole. We therefore asked whether reducing Nkx2-5 activity could restore regular heartbeat in the absence of Isl1. Efficient suppression of Nkx2-5 was achieved using validated CRISPR/Cas9-mediated F0 mutagenesis (Figure S5). Consistent with previous studies, *nkx2-5* crispants displayed regular cardiac rhythm despite expansion of the pacemaker domain (Figure 5A, G). We next simultaneously disrupted *isl1* and *nkx2-5* and analyzed cardiac electrophysiology using VSFP imaging at 3 dpf. Remarkably, loss of *nkx2-5* restored the regular generation of action potentials in *isl1*-deficient embryos (Figure 5A, compare with Figure1B). Atrial and ventricular repolarization kinetics (APD50), which were altered in *isl1* mutants, returned to values comparable to wild type (Figure 5B, C). Likewise, conduction velocity remained unaffected (Figure 5D) and beat-to-beat variability was restored (Figure 5E), demonstrating rescue of the electrophysiological defects caused by loss of Isl1.

**Figure 5:**
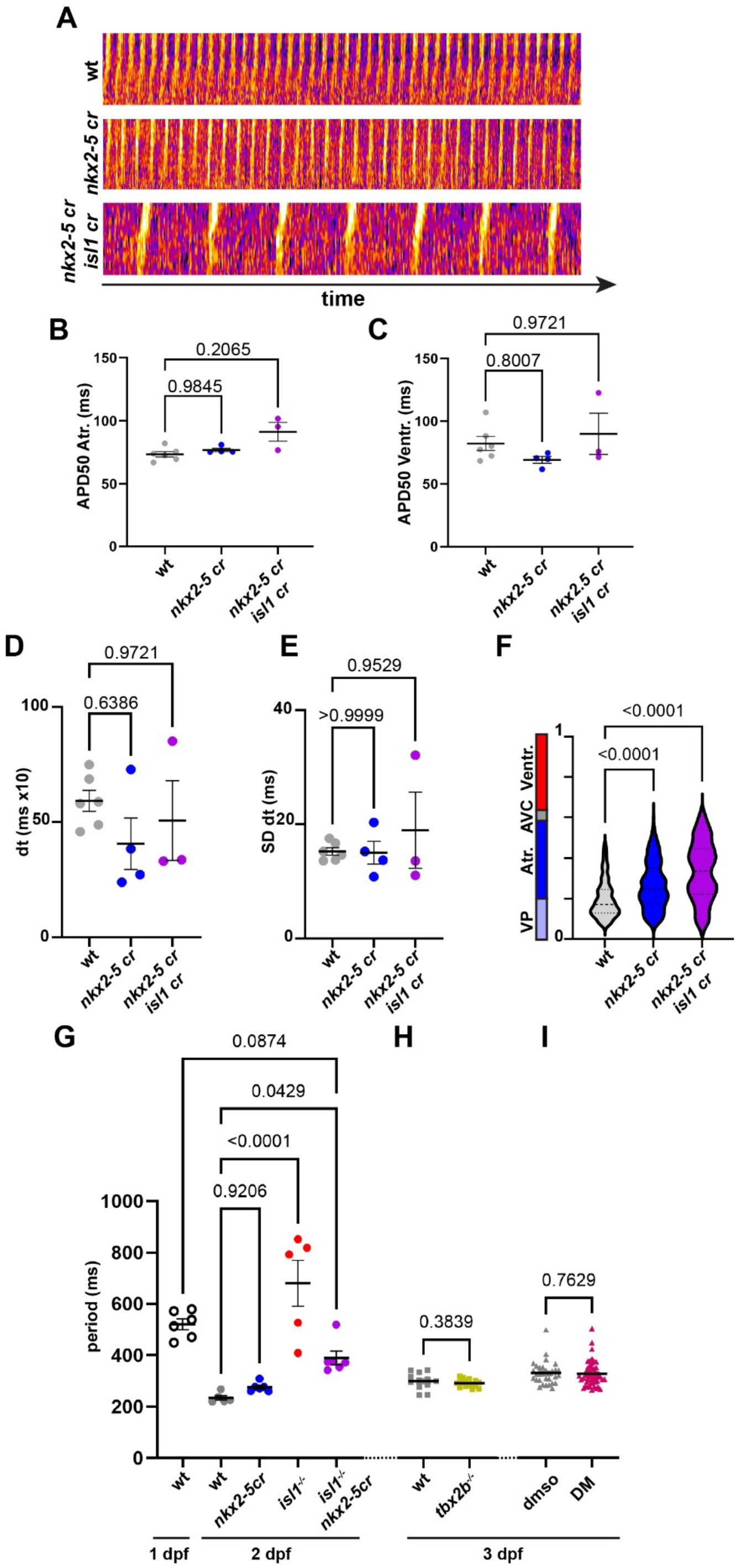
Suppression of *nkx2-5* expression rescues AP generation and heartbeat regularity in *isl1* defective embryos. **(A)** Ratiometric profiles of fluorescent intensities of APs recorded in 3 dpf *tg(myl7:VSFP-CY-Butterfly)* crispants for wt, *nkx2-5* only and combination of *nkx2-5* and *isl1.* When both genes are targeted, rescue of the regularity, albeit at a slower of generation rate, of the APs is achieved (compare with Figure 1B). **(B)** and **(C)** Atrial (B) and ventricular (C) APD50 values for *nkx2-5* and *nkx2-5;isl1* defective larvae are not significantly different from wt. **(D)** Duration of the propagation of the AP across embryonic heart segments happens is maintained in *nkx2-5* and *nkx2-5;isl1* crispant embryos. **(E)** AP generation regularity is rescued by suppression of *nkx2-5* expression in Isl1-defective embryos. **(F)** Mapping of AP points of origin reveals that in the absence of *nkx2-5*, the pacemaker region expands within the atrium and apparently beyond when both Isl1 and Nkx2-5 expression are perturbed. **(G)** Regularity of the heartbeat contraction period is rescued in *isl1^-/-^* embryos crispant for *nkx2-5.* Note that their heartbeat period is not significantly different from that of a 1dpf embryonic wt heart. **(H)** Genetic suppression of Tbx2b expression or **(I)** chemical suppression of bmp activity by dorsomorphin (DM) treatment do not affect pacemaker function. Panels B-G: one-way ANOVA with multiple comparisons, wt values taken as reference. Bars are at mean +/-SEM. Panels H,I: Two-tailed, non-paired Student’s t-test; p value is reported in the figure panel. Bars are at mean +/-SEM.

Mapping the origin of individual action potentials revealed that *nkx2-5* crispants initiate action potentials over a broader region of the atrium than wild-type embryos (Figure 5F), consistent with expansion of the pacemaker program previously reported in *nkx2-5* mutants both in mouse (Mommersteeg et al., 2007) and in zebrafish (Colombo et al., 2018). Double *isl1;nkx2-5* crispants showed a similarly broad distribution of action potential initiation sites, but unlike *isl1* mutants these initiation events occurred regularly and produced a stable cardiac rhythm (Figure 5G), reflecting the intrinsic automaticity of embryonic cardiomyocytes, which is not affected by loss of *isl1*. Interestingly, the slower heart rate observed in double *isl1;nkx2-5* crispants resembled that of wild-type embryos at 24 hpf, before the dominant pacemaker becomes established.

We next asked whether the observed restored regular pacing was accompanied by recovery of the Isl1-dependent transcriptional program. Unexpectedly, expression of *bmp4* and *tbx2b* remained absent from the venous pole of double *isl1;nkx2-5* crispants (Figure S5D). Furthermore, neither suppression of *tbx2*b nor pharmacological inhibition of BMP signaling affected cardiac rhythm (Figure 5H, I), indicating that these Isl1 downstream targets are not required for embryonic pacemaker activity.

Altogether, these findings demonstrate that ectopic Nkx2-5 activity underlies the electrophysiological defects caused by loss of Isl1. Notably, restoration of regular pacing occurs despite the continued absence of *bmp4* and *tbx2b* expression, genetically separating Isl1-mediated repression of the chamber myocardial program from its regulation of other pacemaker-associated genes. These results identify abnormal Nkx2-5 activity, rather than loss of *bmp4* or *tbx2b* expression, as the principal cause of the pacemaker defects in Isl1-deficient embryos.

## Discussion

In this study, we combined high-speed optical mapping with genetic perturbation to resolve the transcriptional hierarchy that establishes and maintains pacemaker identity in the zebrafish embryonic heart. We identify Isl1 as a central regulator of this process. Loss of Isl1 does not prevent the incorporation of venous pole cardiomyocytes but instead causes these cells to lose pacemaker identity and acquire characteristics of chamber myocardium through ectopic *nkx2-5* expression. Reducing Nkx2-5 activity largely restores regular rhythm in Isl1-deficient embryos despite the persistent absence of the Isl1-dependent pacemaker gene program. This suggests that reducing Nkx2-5 activity unmasks the intrinsic, Isl1-independent automaticity of the embryonic heart tube, allowing regular activation from a broader posterior region of the heart in the absence of the dominant pacemaker localized at the venous pole.

Together, our findings reveal that during zebrafish heart development the reciprocal inhibition of Isl1 and Nkx2-5 establishes a boundary between pacemaker and working myocardium at the venous pole. While the boundary-defining role of Nkx2-5 is conserved, the factor that restricts it within the pacemaker domain is not: this function is performed by Isl1 in zebrafish and by SHOX2 in mouse.

Recent work has substantially advanced our understanding of early pacemaker development in zebrafish. Antagonistic BMP and Hedgehog signaling specifies the inflow tract progenitor population during gastrulation, after which Wnt signaling promotes pacemaker differentiation (Robertson et al., 2026). These signaling pathways therefore determine the formation and size of the pacemaker lineage before myocardial differentiation. Our findings position the Isl1-Nkx2-5 regulatory network downstream of these early patterning events. Consistent with previous work showing that venous pole cardiomyocytes are incorporated normally in *isl1* mutants (Burkhard et al., 2017), Isl1 is not required for specification of pacemaker progenitors but instead acts after their differentiation to preserve pacemaker identity by preventing activation of the chamber myocardial program. Together, these studies suggest that zebrafish pacemaker development proceeds through successive regulatory modules that first specify, then differentiate and finally stabilize the pacemaker lineage.

In mammals, the SAN and chamber myocardium arise from a common pool of second heart field progenitors. These lineages are distinguished by their Nkx2-5 expression: Tbx18-, Shox2-, Tbx3- and Isl1-positive cells destined for the SAN actively repress Nkx2-5, whereas chamber progenitors activate Nkx2-5 and suppress the pacemaker program (Blaschke et al., 2007; Christoffels et al., 2006; Espinoza-Lewis et al., 2009; van der Maarel and Christoffels, 2024; Mohan et al., 2018; Mommersteeg et al., 2007). Isl1 is important for the specification and expansion of second heart field derivatives, while its sustained expression within the developing SAN depends on Shox2-mediated repression of Nkx2-5. Our findings demonstrate that this hierarchy does not apply to the zebrafish embryonic heart. Venous pole cardiomyocytes are incorporated normally in zebrafish *isl1* mutants (Burkhard et al., 2017), placing Isl1 downstream of progenitor specification and specifically at the level of the maintenance of pacemaker identity. Furthermore, *isl1* expression is maintained in *shox2* mutants, whereas *shox2* expression is lost in *isl1* mutants, placing Isl1 upstream of Shox2 in the zebrafish gene regulatory network.

These conclusions differ from previous morpholino-based studies that proposed an essential role for Shox2 in zebrafish pacemaker development. Although we cannot formally exclude residual Shox2 activity in the three independently generated mutant alleles analyzed here, all three yielded concordant phenotypes, with preserved *isl1* expression, maintenance of the Nkx2-5-negative pacemaker domain, and normal embryonic cardiac rhythm. This contrasts with the mouse, where SHOX2 is an essential suppressor of NKX2-5 in the SAN (Espinoza-Lewis et al., 2009) and *Nkx2-5* inactivation in the Shox2 domain rescues pacemaker cell fate (Li et al., 2024). In zebrafish, this role is not fulfilled by Shox2, but by Isl1 instead.

In mammals, the *Isl1*-dependent second heart field contributes extensively to multiple cardiac structures, requiring lineage-specific gene regulatory networks to distinguish the SAN cells from the chamber myocardium. In this context SHOX2 is required to exclude NKX2-5 specifically within the SAN lineage and may overshadow the function of ISL1 in the SAN. In contrast, the zebrafish second heart field contributes comparatively few cells to the embryonic heart, including the venous pole. We therefore propose that Isl1 itself continuously prevents Nkx2-5 expression within the venous pole, eliminating the need for an intermediate regulator such as Shox2. This model may explain why Isl1 loss causes expansion of Nkx2-5 into an already established pacemaker domain rather than a failure of venous pole formation, and why Shox2 becomes dispensable for embryonic pacemaker development in zebrafish.

An important conceptual finding of this study is that restoration of regular rhythm can be genetically uncoupled from complete restoration of the Isl1-dependent gene regulatory network. Reducing Nkx2-5 activity largely normalized action potential initiation and rhythm in Isl1-deficient embryos, yet expression of *bmp4* and *tbx2b* remained absent and heart rate was not fully restored. These observations indicate that Isl1 regulates multiple downstream pathways; on one hand Isl1 restricts *nkx2-5* expression and thereby suppresses the chamber myocardial differentiation program, which blocks automaticity. This is illustrated by the fact that upon Nkx2-5 reduction, pacemaker activity shifts to the entire posterior segment of the heart tube. On the other hand, Isl1 regulates a pacemaker-specific gene program, including *bmp4, tbx2b and hcn4*, that may contribute to maturation, robustness, or fine-tuning of pacemaker function.

Our genetic data further suggest that reciprocal repression between Isl1 and Nkx2-5 forms a bistable transcriptional switch that stabilizes the boundary between pacemaker and chamber myocardium. Reciprocal transcriptional repression is a recurring developmental strategy for generating stable boundaries between adjacent cell populations with distinct identities. A classical example is the mutual repression between Pax2 and Pax6, which establishes and maintains the optic stalk-retina boundary during eye development (Schwarz et al., 2000). We propose that the Isl1-Nkx2-5 network fulfills an analogous role in the zebrafish embryonic heart. Unlike many developmental boundaries that become irrelevant after organogenesis, the interface between pacemaker and chamber myocardium must remain stable throughout life while maintaining direct electrical coupling between tissues with fundamentally different electrophysiological properties. A mutually antagonistic Isl1-Nkx2-5 circuit provides an elegant mechanism to preserve this interface by simultaneously reinforcing pacemaker identity and suppressing acquisition of chamber characteristics. The molecular mechanism underlying the antagonistic relationship between Isl1 and Nkx2-5 remains incompletely understood. NKX2-5 and ISL1 expression has been shown to overlap in a subset of mammalian cardiac progenitors (Ma et al., 2008). NKX2-5 can directly repress ISL1 through binding of conserved enhancer elements and the ISL1 promoter (Dorn et al., 2015), and NKX2-5 deficiency increases ISL1 expression in human iPSC-derived cardiac progenitors (Anderson et al., 2018). However, there is so far no evidence that ISL1 directly represses NKX2-5, but actually more the existence of extensive co-occupancy of ISL1 and NKX2-5 at shared enhancers (Maven et al., 2023) where the two transcription factors may compete, resulting in transcriptional profiles that reflect their relative abundancy (Watanabe et al., 2012). Our own genetic data and previous observations by Colombo et al., 2018 support a model in which reciprocal antagonism between Nkx2-5 and Isl1 stabilizes the boundary between pacemaker and chamber myocardium during zebrafish heart development. Determining whether this interaction involves direct transcriptional repression will require future genomic and biochemical studies.

Our data further distinguish the role of Isl1 from more general regulators of cardiac electrophysiology. Whereas Tbx5a deficiency slowed atrioventricular conduction and prolonged repolarization in both cardiac chambers, consistent with its broad role in myocardial differentiation and electrical maturation, Isl1 deficiency had relatively modest effects on conduction velocity and repolarization but abolished the dominance of the venous pole as the site of action potential initiation. This functional dissociation demonstrates that Isl1 acts specifically to preserve pacemaker identity rather than to regulate myocardial electrophysiology more generally, and that the bradyarrhythmia in *isl1* mutants results directly from molecular reprogramming of venous pole cardiomyocytes towards a chamber-like identity.

Together, our findings establish a zebrafish-specific gene regulatory network in which Isl1 safeguards pacemaker identity through continues restriction of Nkx2-5 independently of Shox2. More broadly, our work demonstrates that highly conserved developmental outcomes can be achieved through distinct transcriptional hierarchies in different vertebrate species. We propose that the Isl1-Nkx2-5 antagonistic circuit represents a core stabilizing module that preserves the boundary between pacemaker and chamber myocardium after its initial specification. Defining how this module interacts with earlier morphogen signaling and with downstream effectors of pacemaker differentiation will provide a more complete understanding of cardiac conduction system development and may ultimately facilitate strategies to generate and maintain authentic pacemaker cells for regenerative medicine.

## Materials and Methods

### Zebrafish Lines

All animal experiments were conducted under the guidelines of the animal welfare committee of the Royal Netherlands Academy of Arts and Sciences (KNAW). Adult zebrafish (Danio rerio) were maintained and embryos raised and staged as previously described (Alestrom et al., 2020) (Westerfield, 2007)

The zebrafish lines used in this study are Tübingen longfin (wild type), *tg(myl7:chimeric VSFP-butterfly CY)* (van Opbergen et al., 2018), *tg(myl7:GFP)* (Huang et al., 2003), *isl1_K88X (isl1 ^sa0029^)* (de Pater et al., 2009), *oug/tbx5a* (Tessadori et al., 2021) and fby/tbx2b (Snelson et al., 2008). Genotyping was carried out as described in the relevant references.

### Generation of the shox2_RL, shox2_HD and shox2_TR alleles

To generate shox2_RL (crRNA9 and crRNA22), shox2_HD (crRNA3 and crRNA71) shox2_TR (crRNA71), Alt-R crRNAs (see Table 1 for sequences) were designed in CHOPCHOP (https://chopchop.cbu.uib.no/; danRer11/GRCz11) and ordered together with tracrRNA at IDT (Leuven, Belgium). RNP complexes with Cas9 (12,5 µg/µL, Hubrecht protein facility) were generated essentially following IDT protocol for Zebrafish embryo microinjection. Microinjections in 1-cell zebrafish embryos (F0), selection of founders, generation and selection of F1 carriers were carried as described previously (Tessadori et al., 2018) with use of the appropriate primers (see Table 1 for primer sequences).

**Table 1:**
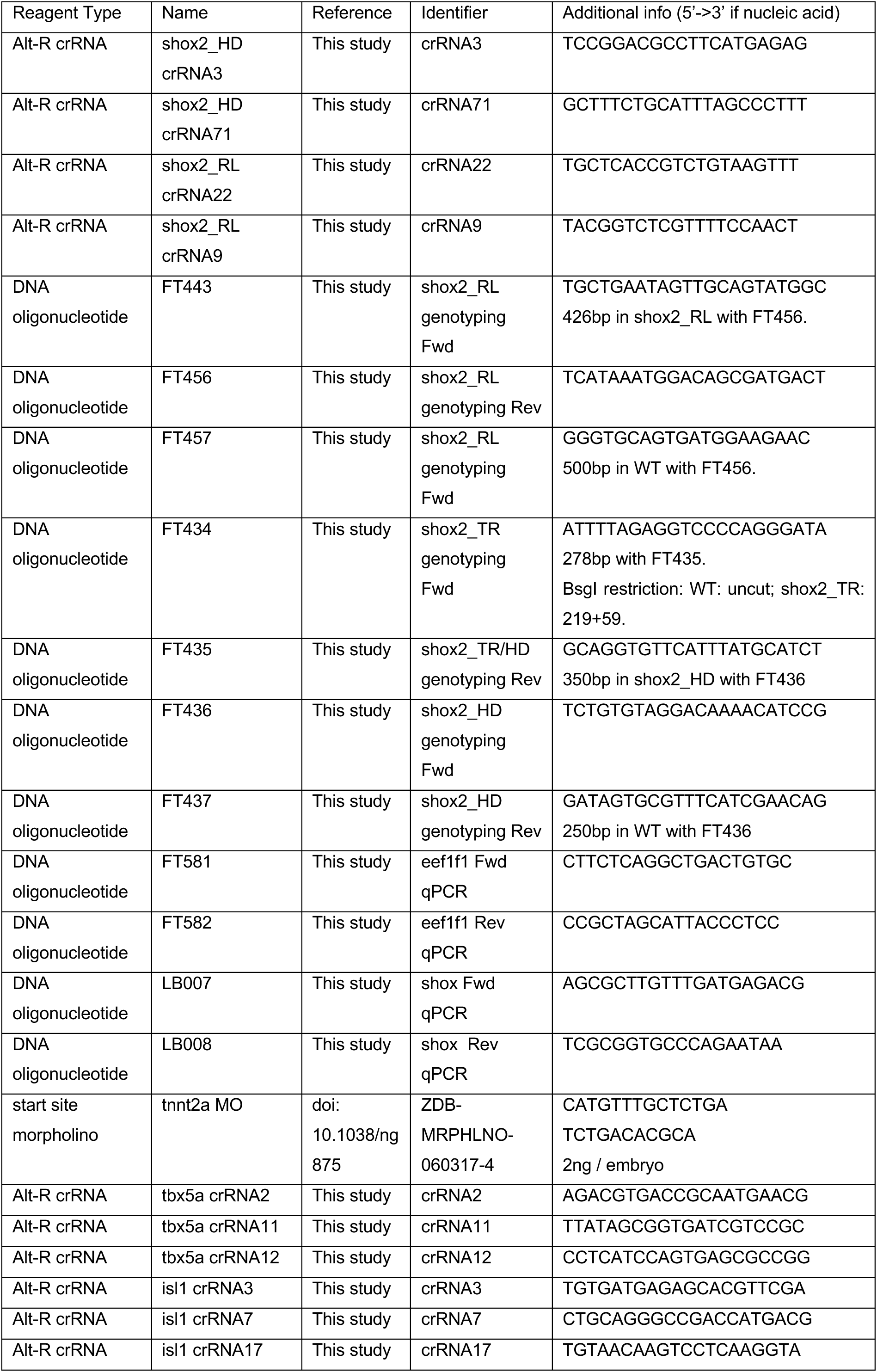
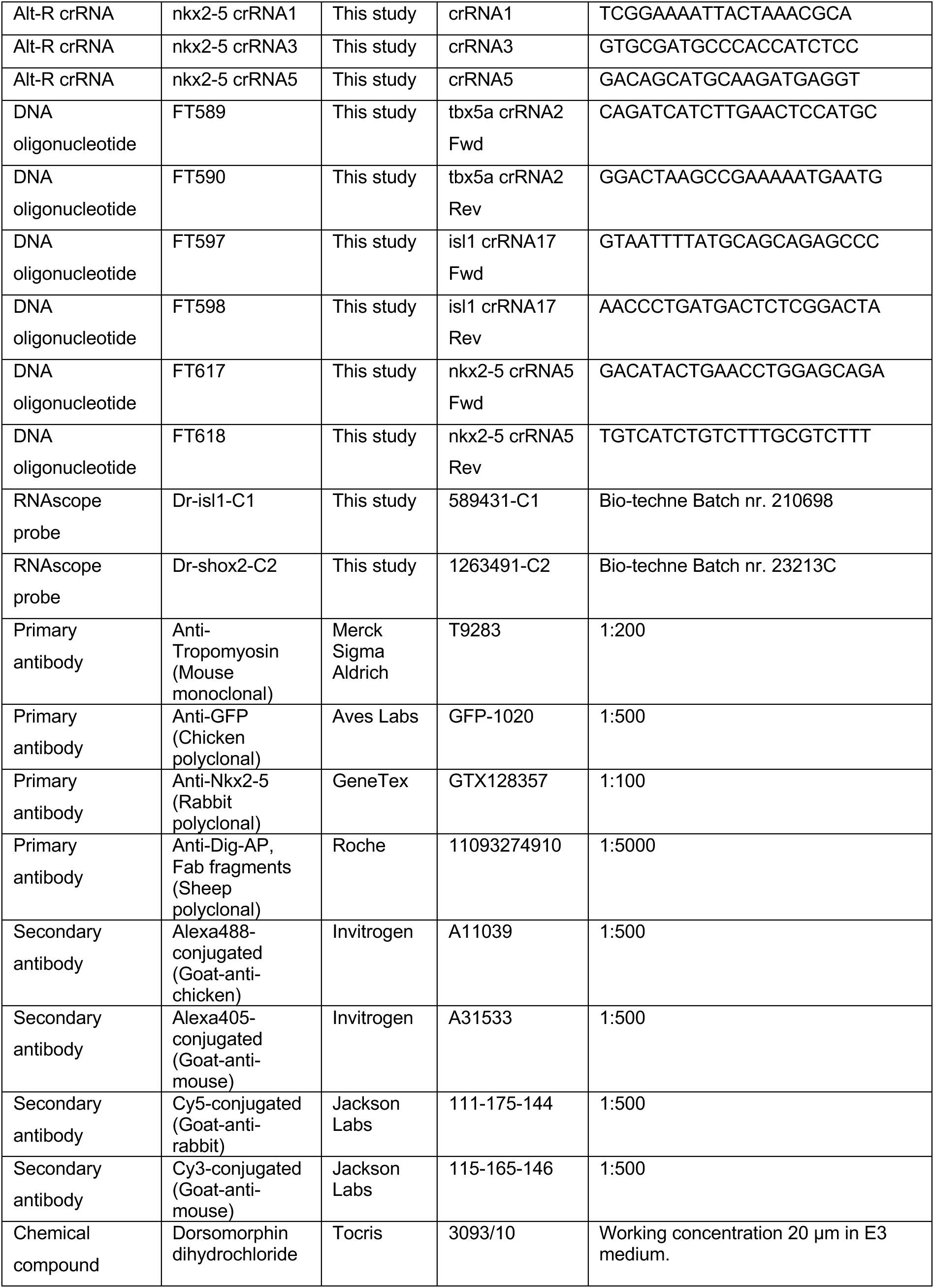
Reagents used in this study.

### Stopping cardiac contraction

To stop cardiac contraction in VSFP experiments, approximately 2ng of a morpholino oligonucleotide targeting the translation start site of *tnnt2a* (Sehnert et al., 2002); see Table 1 for sequence) was injected in one-cell stage embryos.

### F0 CrispR/Cas9 injections (*Crispants* generation)

To generate F0 embryos displaying loss-of-function phenotypes for *isl1, tbx5a* or *nkx2-5* or any combination of these, design, RNP generation and microinjection in one-cell embryos was carried out essentially as for the generation of shox2 alleles (see above), with as main difference that for each targeted gene 3 different RNPs were injected simultaneously. Details of the targeting position for each RNP can be found in the corresponding supplementary figures. Sequences of the used guides are reported in Table 1. Prior to inclusion in any experiment the embryos were checked for exclusive presence of the expected phenotype as displayed by the genetic mutant.

### Whole mount mRNA In Situ Hybridization (ISH)

Embryos were fixed 4% paraformaldehyde (PFA) either overnight at 4°C or for 2 hours at room temperature (RT) and stored in methanol (MeOH) at −20°C. Rehydration of the embryos was carried out in PBST (PBS, 0.1% Tween-20) and embryos were treated with Proteinase K (1 µg/ml, Promega) for 10 min at RT. After rinsing in PBST, embryos were post-fixed in 4% PFA for 20 min, washed in PBST and pre-hybridized for at least 1 h in Hyb-buffer (recipe). Digoxigenin-labeled were diluted in Hyb-buffer supplemented with transfer RNA (Sigma-Aldrich) and heparin (Sigma-Aldrich), and incubated with the embryos overnight at 70°C. After removal of the probe, embryos were washed stepwise from Hyb- to 2xSSCT, and subsequently from 0.2xSSCT to PBST. Blocking of the samples was carried out for at least 1 h at room temperature (RT) in PBST supplemented with sheep serum and Bovine Serum Albumin (BSA) before incubation overnight at 4°C with an sheep-anti-digoxygenin-AP antibody (1:5000; Cat: 11093274910; Roche). Embryos were then washed in PBST before transfer to TBST and staining with Nitro-blue tetrazolium/5-bromo-4-chloro-3-inodyl phosphate (NBT/BCIP; Cat: 11093274910; Roche) in TBST. Embryos were washed extensively in PBST and fixed overnight in 4% PFA at 4°C, and cleared in MeOH. Imaging was carried out in benzylbenzoate:benzylalcohol (2:1; Murray’s).

Genes assayed by whole-mount ISH: *myl7* (NM_131329), *nppa* (NM_198800), *tbx2b* (NM_131051), *bmp4* (NM_131342), *hcn4* (XM_680986), *shox2* (NM_201196) and *shox* (NM_001126411).

### RNAscope

Whole-mount RNAscope (ACD, Bio-Techne, Minneapolis, MN, USA) was carried out on fixed zebrafish embryos essentially as described in (Gross-Thebing et al., 2014) for sample preparation and probe hybridization and according to the manufacturer’s protocol (ACD technical note on RNAscope on Whole Zebrafish Embryos) for detection steps. Details of the probes used are reported in Table 1. When combined with immunofluorescent labeling (IF, see below), RNAscope was carried out first, directly followed by IF.

### Immunofluorescent labeling

Zebrafish embryos at the appropriate developmental stage were fixed for 2 hours at RT or overnight at 4°C in 4% PFA. After washing with 1×PBS–Triton X-100 (0.1%; PBSTx) and blocking in 10% goat serum in 1×PBSTx (blocking buffer; BB), embryos were incubated overnight at 4°C with primary antibodies. Details and dilutions used are reported in Table 1. After washing in PBSTx, the samples were incubated for 2 hours at RT with secondary antibodies in BB. Details and dilutions used are reported in Table 1. Embryos were washed in PBSTx prior to mounting and imaging.

### Quantitative RT-PCR

Total RNA and genomic DNA extraction was performed with TRIZOL (LifeTechnologies/Thermo Fisher Scientific, Bleiswijk, the Netherlands) on single 2 dpf zebrafish embryos from an incross of *shox2_RL, shox2_TR or shox2_HD* heterozygous carriers, following the manufacturer’s protocol. Genomic DNA was used to determine the genotype of the single embryos (see Table 1 for primer sequences). cDNA was obtained with RT Superscript III (Life Technologies/Thermo Fisher Scientific) and used as template for qPCR using a CFX Connect96 Real-Time System (Bio-Rad Laboratories, Hercules, CA, USA) and SYBR green qPCR Master Mix (Thermo Fisher Scientific, Waltham, MA, USA). Primers used for assessing *shox* expression are reported in Table 1. Relative gene expression (2-ΔΔCt) was calculated after normalization of *shox* expression to the average expression observed in WT single embryos using *eef1g* as a reference.

### Imaging

#### Live embryos

Assessment of *in vivo* of phenotypes was carried out on a Leica M165FC stereomicroscope or a Zeiss StemiSV6 stereomicroscope (Carl Zeiss AG, Oberkochen, Germany).

High-speed heartbeat recordings were carried out at 100 frames per second (fps) on a Leica DMI6000 SD microscope (Leica Microsystems, Wetzlar, Germany) with a Hamamatsu digital camera C13440 (Hamamatsu Photonics, Hamamatsu, Japan) in a 28.5°C controlled chamber. BMP inhibition was achieved by treating embryos between 28 hpf and time of recording with 20µm dorsomorphin dihydrochloride in E3.

VSFP recordings were carried out essentially as described in van Opbergen et al., 2018. Images were collected using a custom-built upright widefield microscope (Cairn Research, Kent, UK) equipped with a 20x 1.0 NA objective (Olympus XLUMPLFLN20X W). White LED excitation light was filtered using a 438/24 nm filter (Semrock FF02-438/24-25) and reflected towards the objective using a 458 nm dichroic mirror (Semrock FF458-Di02-25×36). Emitted fluorescence was directed to an emission splitter unit (OptoSplit II ByPass ImageSplitter) fitted with a 509 nm dichroic mirror (Semrock FF509-FDi01-25×36) and 483/32 nm and 514 long-passemission filters (Semrock FF01-483/32-25 and LP02-514RU-25, respectively). Images were projected on a high-speed camera (Andor Zyla 4.2 plus sCMOS). Recordings were performed at 100 fps, generally for 500-1500 frames.

For all live imaging, sedation of embryos was achieved with 16,8 mg/ml tricaine (MS222; Sigma-Aldrich) in E3 medium. Embryos were mounted in 0,3% Agarose in E3 embryo water supplemented with sedative.

#### Fixed samples

Whole-mount ISH samples were imaged on a Zeiss Axioplan microscope (Carl Zeiss AG) with a DFC420 digital microscope camera (Leica Microsystems).

Confocal imaging was carried out on a Leica SP8 confocal microscope (Leica Microsystems).

### Image Analysis

#### Live embryos

Ratiometric analysis of VSFP recordings and obtention of AP profiles was carried out using ImageJ/Fiji (https://imagej.net/software/fiji/).

Calculation of APD50 values was carried out in Matlab (Version R2024b, Mathworks, Natick, MA, USA).

Action potential characterization and quantification were performed using a simple script run in R (The R foundation, https://www.r-project.org/) that reads the pixel values and pixel coordinates of the AP profiles obtained from the VSFP recordings. For each kymograph, the script runs through the frames in chronological order until it finds enough high intensity pixels to qualify as the start of the action potential. It marks the end of the action potential once the number of high intensity pixels falls below the threshold and records all values in between. From the recorded spatial and temporal limits of the action potentials, means, medians, and standard deviations were produced. These were then used to filter out false positives and noise.

### Fixed Samples

#### RNAscope assessment

Expression levels of *shox2* and *isl1* in the heart were quantified in embryos in which *shox2* and *isl1* transcripts were detected by RNAscope and cardiomyocytes (CMs) were labeled with Tropomyosin or GFP, depending on the presence of a *myl7* transgene. In Imaris (Oxford Instruments, Abingdon, United Kingdom), the heart was first masked using the CM-specific labeling, removing the background signal. The specific signal for *isl1* and *shox2* was then measured using the Spots feature in Imaris. Automatic detection by Imaris was inspected and confirmed or false positives were manually removed. Quantifications were carried out blindly to the genotype of the embryos.

#### Nkx2-5-negative myocardium quantification

Hearts labeled by immunofluorescence for Nkx2-5 and GFP (*tg(myl7:GFP);* CM) were used to measure the volume of Nkx2-5-atrial myocardium, defined by CM+ Nkx2-5-labeling. Confocal z-stacks were processed in Imaris software (Oxford Instruments, Abingdon, United Kingdom). After using a CM marker mask, 2 dpf hearts were clipped at the atrioventricular canal and at the boundary of the Nkx2-5 expression domain at the VP, thereby defining the Nkx2-5+ atrial myocardium volume. The same procedure was carried out on the entire atrium, resulting in the total atrial volume. Subsequently, the Nkx2-5-volume is obtained and its ratio (expressed as percentage) on the total atrial volume is quantified.

#### Statistics

All statistical assays were carried out in Graphpad Prism 9.0 or higher (GraphPad Software). Used tests and significance levels are reported in figures and corresponding legends.

## Acknowledgements

The authors wish to thank Anko de Graaff and Pim Toonen (Hubrecht Imaging Center) for technical assistance, the Hubrecht Animal Facility for maintenance of the zebrafish, Hiroyasu Kamei and Shin-Ichiro Takahashi (Kanazawa University, Kanazawa, Japan) for sharing the *shox* coding sequence and Vincent Christoffels for fruitful discussions throughout the project and critically reading the manuscript. This work was supported by NWO grant OCENW.GROOT.2019.029 (F.T & J.B.). Work by A.K. reported in this study was part of an internship supported by the Hubrecht Talent Program.

---

**Supplementary Figure SF1 related to Figure 1:**
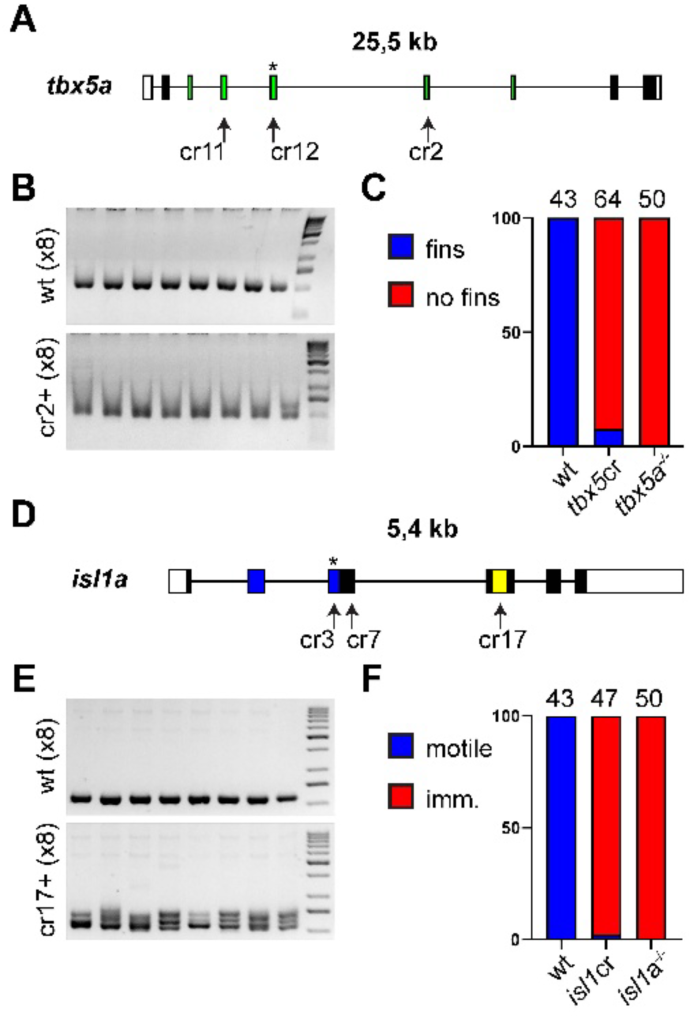
F0 crispants for *tbx5a* and *isl1*: strategy, confirmation and phenotyping. **(A)** Genomic locus of *tbx5a*. The crRNAs were designed to target different exons encoding the homeodomain (green). Asterisk denotes the position of the point mutation resulting in the *tbx5a/oug* null allele (Tessadori et al., 2021). **(B)** Genome editing is confirmed by AFLP-based PCR on crRNA sequence target (only cr2 is shown, other crRNAs show comparable results). While uninjected control embryos display sharp 217bp PCR products, their smeary appearance in injected embryos (lower panel) denotes successful genome editing. **(C)** Phenotyping of F0 crispants and comparison with genetic allele *tbx5a/oug* confirms efficiency and validity of approach. **(D)** Genomic locus of *isl1*. The crRNAs were designed to target different exons encoding the LIM domain (blue) and homeodomain (yellow). Asterisk denotes the position of the point mutation resulting in the *isl1K88X* null allele used in this study (De Pater et al., 2009). **(E)** Genome editing is confirmed by PCR on crRNA sequence target (only cr17 is shown, other crRNAs show comparable results; PCR product in control embryos is 217bp). **(C)** Phenotyping of F0 crispants and comparison with *isl1K88X* allele confirms efficiency and validity of approach. For panels C and F number of embryos phenotyped is indicated above histograms.

**Supplementary Figure SF3 related to Figure 3:**
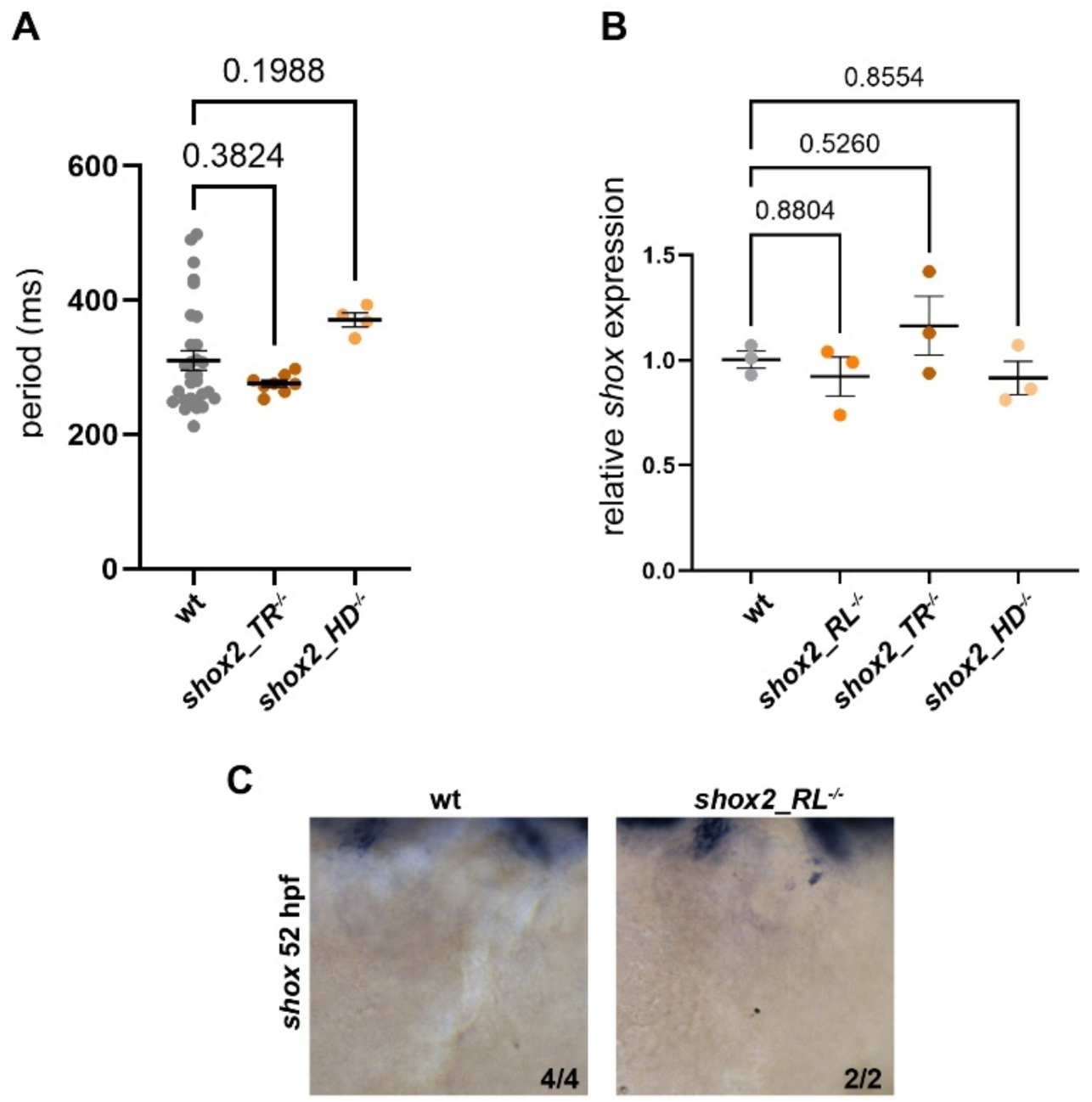
Novel alleles of *shox2* do not elicit genetic compensation by *shox* and show no heartbeat phenotype. **(A)** Heartbeat of *shox2_TR* and *shox2_HD* null embryos is not affected. **(B)** Levels of expression of *shox* are similar between wild type and all three *shox2* alleles presented in this study. **(C)** Expression of *shox* in the wt and *shox2_RL^-/-^* embryonic hearts at 52 hpf. *shox* expression is not detected in the VP in both genotypes. Panels A and B: one-way ANOVA with multiple comparisons, wt values taken as reference. Bars are at mean +/-SEM.

**Supplementary Figure SF4 related to Figure 4:**
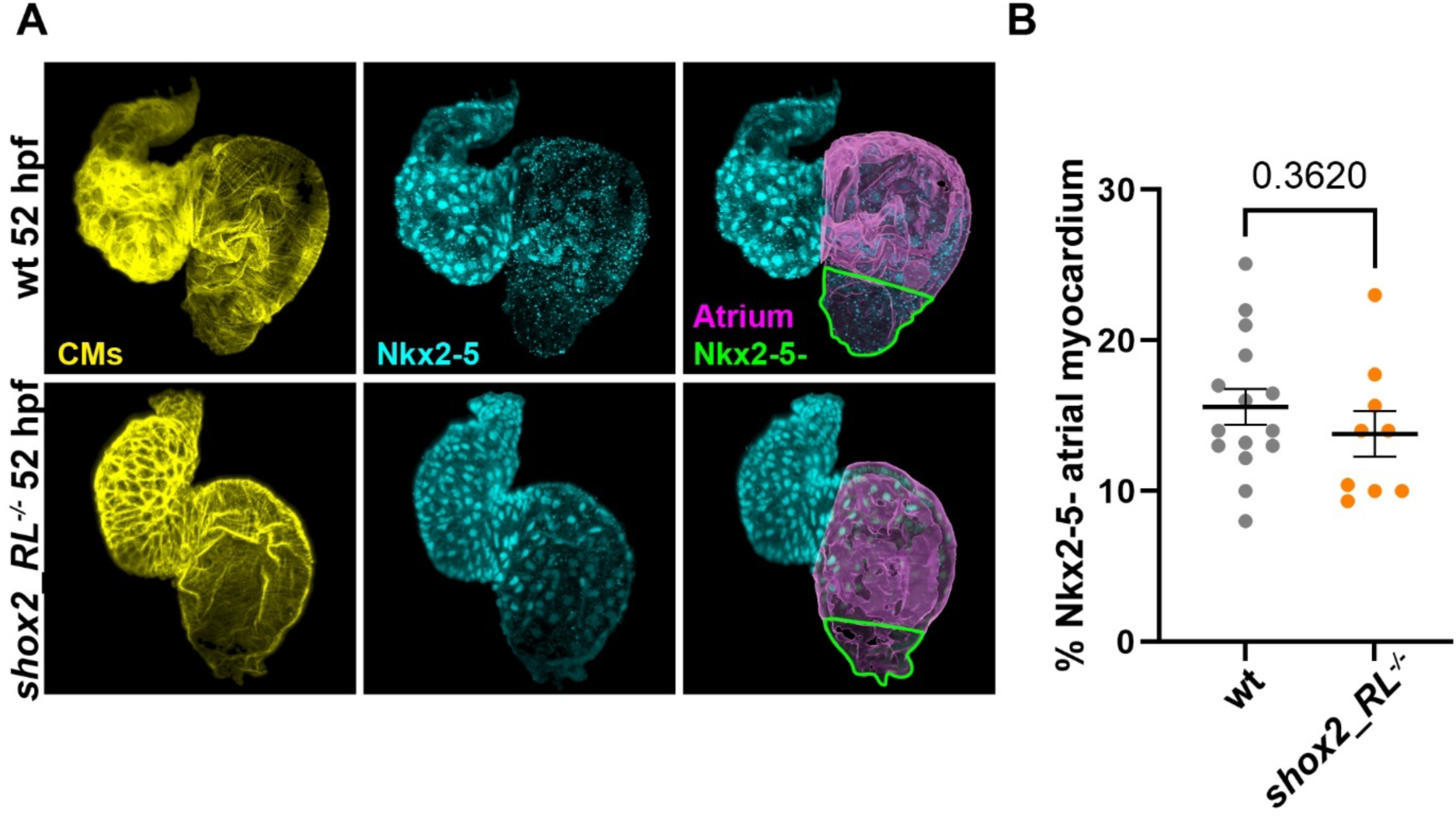
Loss of Shox2 does not affect the boundary of the Nkx2-5 expression domain in the embryonic zebrafish heart. **(A)** Fluorescent immunolabeling of 52 hpf CMs (yellow) for Nkx2-5 (cyan). Atria are overlayed in magenta and the Nkx2-5-VP is outlined in green. **(B)** Quantification of the Nkx2-5-atrial myocardium obtained on confocal imaging as presented in panel **(A)**. Panel B: Two-tailed, non-paired Student’s t-test; p value is reported in the figure panel. Bars are at mean +/-SEM.

**Supplementary Figure SF5 related to Figure 5:**
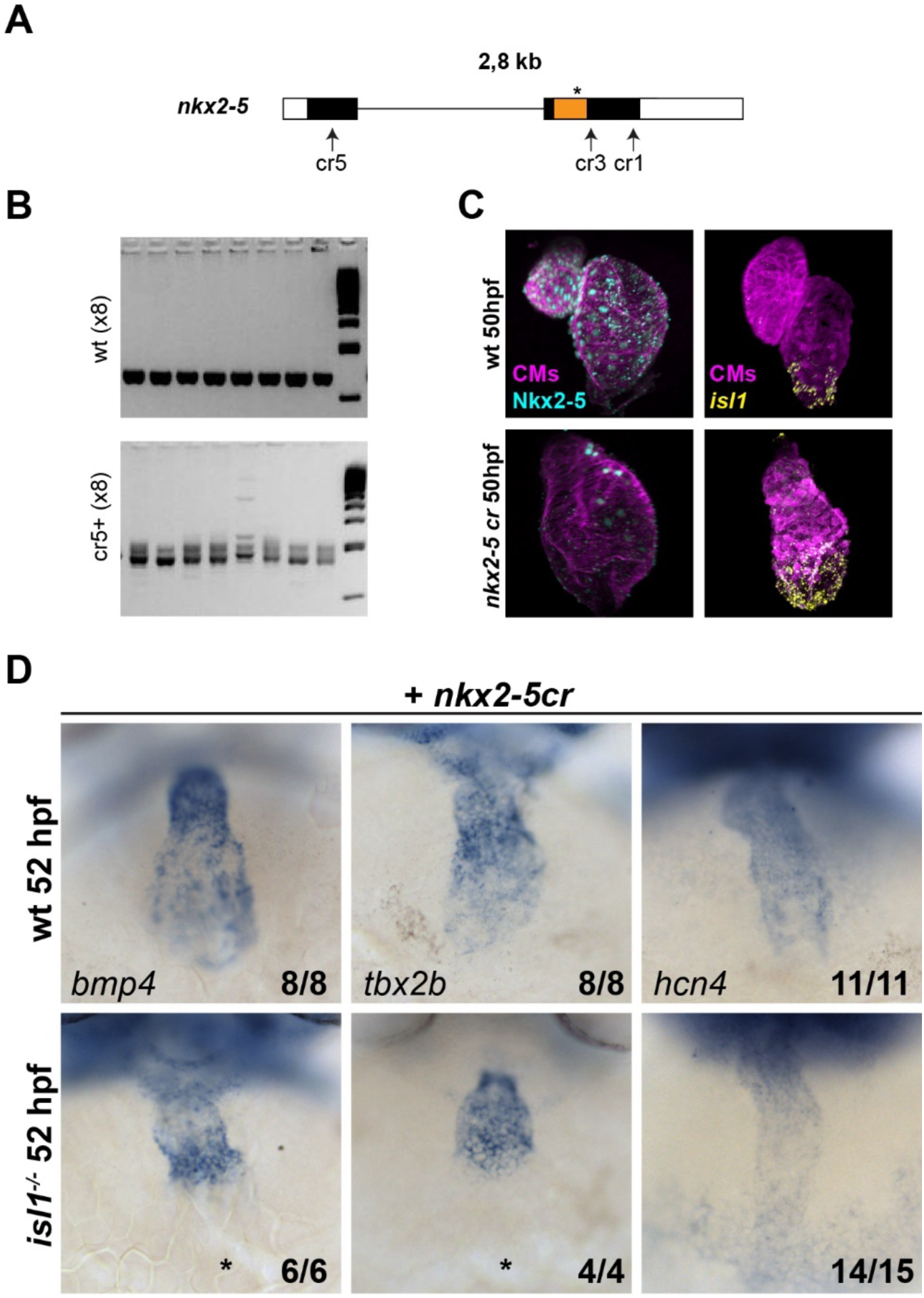
Characterization of the F0 *nkx2-5* crispants. **(A)** Genomic locus of *nkx2-5*. The crRNAs were designed to target different exons encoding the homeodomain (orange). Asterisk denotes the position of the point mutation resulting in the null allele (Targoff et al., 2013). **(B)** Genome editing is confirmed by PCR on crRNA sequence target (only cr21 is shown, other crRNAs show comparable results; PCR product is 288bp). Observation of smears in injected embryos (lower panel) when compared to control (upper panel) denotes successful genome editing. **(C)** Immunolabeling of cardiomyocytes (CMs; Tropomyosin, magenta) and Nkx2-5 (cyan) confirms efficient KO of Nkx2-5 in F0 crispants. Note absence of Nkx2-5 signal and in lower panel typical cardiac phenotype due to Nkx2-5 deficiency (Targoff et al., 2013). Concomitantly, *isl1* expression domain is expanded as detected by RNAscope (yellow). **(D)** Crispant Nkx2-5 loss-of-function embryos display expansion of expression of *bmp4, tbx2b* and *hcn4* in both heart chambers (upper panels). In *nkx2-5* crispants in the *isl1^-/-^* background (lower panels) expression of *bmp4* and *tbx2b* is limited to the AVC and ectopic expression in the ventricle but is absent from the VP (asterisk). Expression of *hcn4* is visible in the entire heart tube, apparently at a lower level than in single *nkx2-5* crispants.

## Notes

### Competing Interest Statement

The authors have declared no competing interest.

